# Sex-dependent and sex-independent brain resting-state functional connectivity in children with autism spectrum disorder

**DOI:** 10.1101/038026

**Authors:** Xin Di, Bharat B. Biswal

**Affiliations:** Department of Biomedical Engineering, New Jersey Institute of Technology, Newark, NJ, USA

**Author notes:** Corresponding author: Bharat B. Biswal, PhD 607 Fenster Hall, University Height Newark, NJ, 07102, USA.

**Keywords:** Autism, sex, female, functional connectivity, resting-state, default mode network

## Abstract

**Background:** Males are more likely to suffer from autism spectrum disorder (ASD) than females. As to whether females with ASD have similar brain alterations remain an open question. The current study aimed to examine sex-dependent as well as sex-independent alterations in resting-state functional connectivity in individuals with ASD compared with typically developing (TD) individuals.

**Method:** Resting-state functional MRI data were acquired from the Autism Brain Imaging Data Exchange (ABIDE). Subjects between 6 to 20 years of age were included for analysis. After matching the intelligence quotient between groups for each dataset, and removing subjects due to excessive head motion, the resulting effective sample contained 28 females with ASD, 49 TD females, 129 males with ASD, and 141 TD males, with a two (diagnosis) by two (sex) design. Functional connectivity among 153 regions of interest (ROIs) comprising the whole brain was computed. Two by two analysis of variance was used to identify connectivity that showed diagnosis by sex interaction or main effects of diagnosis.

**Results:** The main effects of diagnosis were found mainly between visual cortex and other brain regions, indicating sex-independent connectivity alterations. We also observed two connections whose connectivity showed diagnosis by sex interaction between the precuneus and medial cerebellum as well as the precunes and dorsal frontal cortex. While males with ASD showed higher connectivity in these connections compared with TD males, females with ASD had lower connectivity than their counterparts.

**Conclusions:** Both sex-dependent and sex-independent functional connectivity alterations are present in ASD.

## 1 Introduction

Autism spectrum disorder (ASD) is more prevalent in males than females (Baxter et al. 2014; Fombonne 1999). Understanding sex differences of neuroanatomical mechanisms underlying ASD can therefore further our knowledge of the etiology and diagnosis of this prevalent neurodevelopmental disorder (Lai et al. 2015). However, due to the unbalanced number of females and males with ASD and limited number of subjects in neuroimaging studies, the numbers of female subjects examined has been very small. Therefore, female subjects have typically been excluded from research.

In recent years, a handful of studies have examined sex differences on brain alterations in ASD by using anatomical (Beacher et al. 2012a; Di and Biswal 2015; Lai et al. 2013; Schaer et al. 2015) and functional MRI data (Beacher et al. 2012b; Schneider et al. 2013). Two studies demonstrated similar larger regional volumes in the temporal lobe in ASD compared with TD in both sexes (Di and Biswal 2015; Lai et al. 2013), indicating sex-independent neuroanatomical mechanisms underlying ASD. While two other studies demonstrated diagnosis by sex interactions in regional gray matter volumes in the right inferior parietal lobe and rolandic operculum (Beacher et al. 2012a) and gyrification in the ventromedial/orbitofrontal cortex (Schaer et al. 2015), which suggest sex-dependent mechanisms underlying ASD. Lastly, two fMRI studies also demonstrated diagnosis by sex interaction in several brain regions during different tasks (Beacher et al. 2012b; Schneider et al. 2013), indicating sex-specific mechanisms.

Although these studies have attempted to localize regional brain alterations in ASD, theories of brain mechanisms underlying ASD usually focused on brain connectivity (Just et al. 2012; Kennedy et al. 2015). The underconnectivity theory attributes ASD to lower anatomical and functional connectivity between the frontal and parietal cortex. Supporting this hypothesis, white matter fractional anisotropy (FA) using diffusion tensor imaging (DTI) was found to be reduced in many brain regions in ASD individuals (Aoki et al. 2013; Barnea-Goraly et al. 2004; Keller et al. 2007). Coherence of resting-state electroencephalogram (EEG) signals in alpha band (8 – 10 Hz) were found to be reduced in ASD individuals (Murias et al. 2007). However, functional connectivity as measured by resting-state fMRI has provided conflicting results. A recent study by Supekar and colleagues has challenged the underconnectivity theory of ASD, and has demonstrated that there are more cases of hyperconnectivity in ASD children (Supekar et al. 2013). Later studies while carefully addressing methodological issues such as head motion, demonstrated that the connectivity patterns are in generally similar in individuals with ASD individuals and TD individuals, and the detectable differences of connectivity is in general reduced in individuals with ASD (Tyszka et al. 2014). Increased connectivity in individuals with ASD was found restrictedly between the basal ganglia and cortex (Cerliani et al. 2015; Di Martino et al. 2014).

To date, only one study examined sex differences in white matter connectivity of the corpus callosum using DTI in preschool age children with ASD (Nordahl et al. 2015). Nordahl et al. (2015) demonstrated mainly sex specific alterations in ASD. For example, a callosal region connected to the orbitofrontal cortex was smaller in males with ASD, and a smaller callosal region connected to the anterior frontal cortex was found in females with ASD. Diffusion properties were not altered in males with ASD in the corpus callosum, but many diffusion properties such as mean diffusivity (MD) were greater in females with ASD. However, there is no study to examine sex differences in resting-state functional connectivity alterations in ASD.

The aim of the present study is to identify sex-dependent or sex-independent functional connectivity alterations in ASD. By leveraging the autism brain imaging data exchange (ABIDE) (Di Martino et al. 2014), we were able to synthesize resting-state fMRI data from multiple sites. Since the age range of most of the sites are between 6 to 20 years old, we only included data whose sample age range fell between 6-20 years old and also included female subjects with ASD. After carefully matching full scale IQ and head motion, we obtained an effective sample of subjects: 28 females with ASD, 49 TD females, 129 males with ASD, and 141 TD males. A 2 (diagnosis) by 2 (sex) design was adopted to study whether females with ASD have different resting-state functional connectivity patterns compared with TD individuals. We computed ROI (regions of interest) based functional connectivity among ROIs that sampled the whole brain (Dosenbach et al. 2010). Next, we performed a 2 (diagnosis) by 2 (sex) analysis of variance (ANOVA) to identify connections that displayed sex dependent connectivity (if the diagnosis by sex interaction was significant) or sex independent connectivity between groups (if the diagnosis by sex interaction was insignificant while the main effect of diagnosis was significant). Because the sex differences on resting-state functional connectivity in ASD are still largely unknown, the current analysis is highly exploratory.

## 2 Materials and Methods

### 2.1 Subjects and inclusion criteria

Resting-state fMRI and structural MRI data were accessed from the Autism Brain Imaging Data Exchange (ABIDE) project (Di Martino et al. 2014). Because most of the datasets in the database were below 20 years of age, we only included datasets that had samples younger than 20 years of age. Secondly, we only incorporated datasets in which the full scale IQ was available. A subject was included only if his or her full scale IQ score was higher than 70. The Leuven_2 sample doesn't have full scale IQ, but has both verbal IQ and performance IQ. The full scale IQ was estimated by taking an average of verbal and performance IQ for each subject. For each remaining dataset, a two (diagnosis) by two (sex) analysis of variance (ANOVA) was used to test whether the IQ scores had significant diagnosis effect and diagnosis by sex interaction. Significant effect of diagnosis on IQ were found in two datasets (Kennedy Krieger Institute and Leuven_2), therefore these datasets were discarded from the analysis. For the remaining datasets, 2 by 2 ANOVA did not show significant main effects or interactions. Thirdly, a subject's data were discarded if the anatomical data failed in our quality control procedure in a separate analysis of the anatomical data (Di and Biswal 2015). Finally, a subject's data were discarded if head motion during the resting-sate scan exceeded certain threshold, which is described below. As a result, six datasets were incorporated in the current analysis: University of Pittsburgh School of Medicine, University of Michigan sample 1, Yale Child Study Center, NYU Langone Medical Center, Stanford University, and University of California Los Angeles sample 1. For each dataset, there were at least two subjects from each of the groups. The effective sample sizes of the four groups were: 28 females with ASD, 49 TD females, 129 males with ASD, and 141 TD males (Details in Table 1).

**Figure 1.**
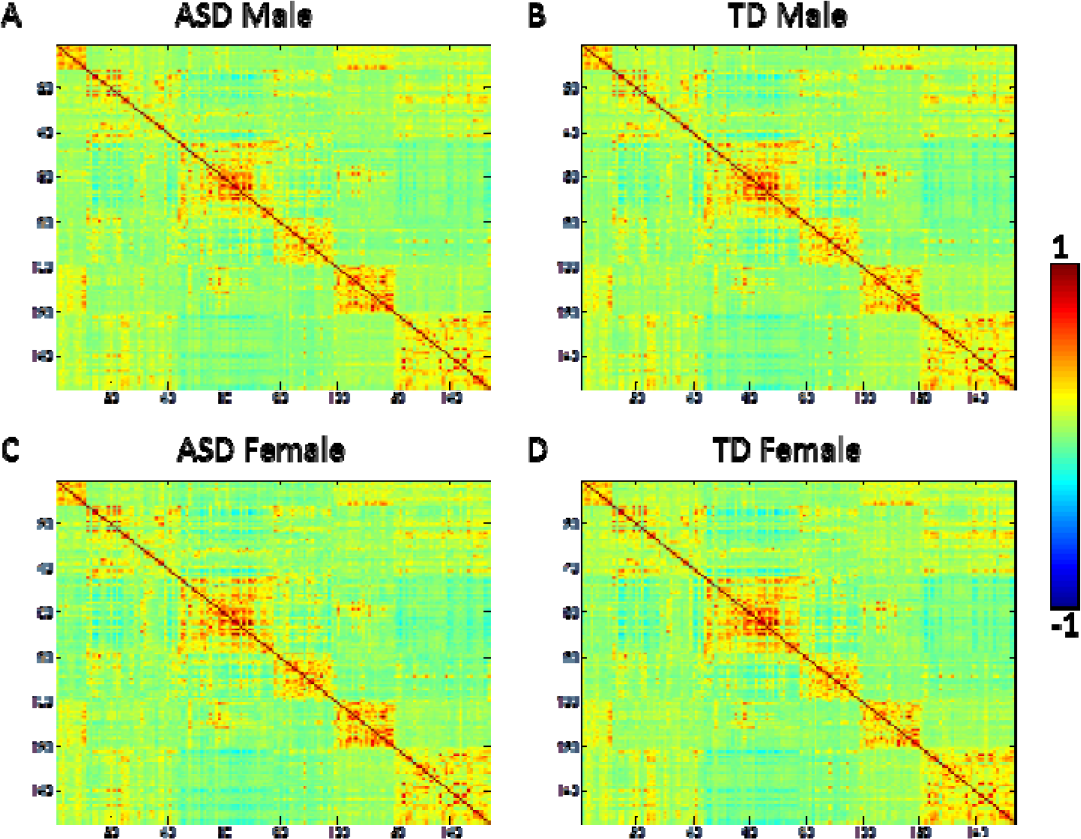
Mean correlation matrices (Fisher’s z score) across 153 regions of interest for the four groups of subjects. ASD, autism spectrum disorder; TD, typically developing.

### 2.2 Resting-state fMRI scanning parameters

All the fMRI and MRI data were scanned using 3T MRI scanners, but with different imaging parameters across sites (Table 2). Subjects were asked to either open or closed their eyes while undergoing an MRI scan in different sites. The repetition time (TR) varied between 1.5 s to 3 s, and the number of time points varied between 120 and 300 (Table 2). In addition to resting-state fMRI images, a high-resolution T1 weighted anatomical images was also acquired for each subject. More details about MRI scanning parameters can be found in (Di Martino et al. 2014).

**Figure 2.**
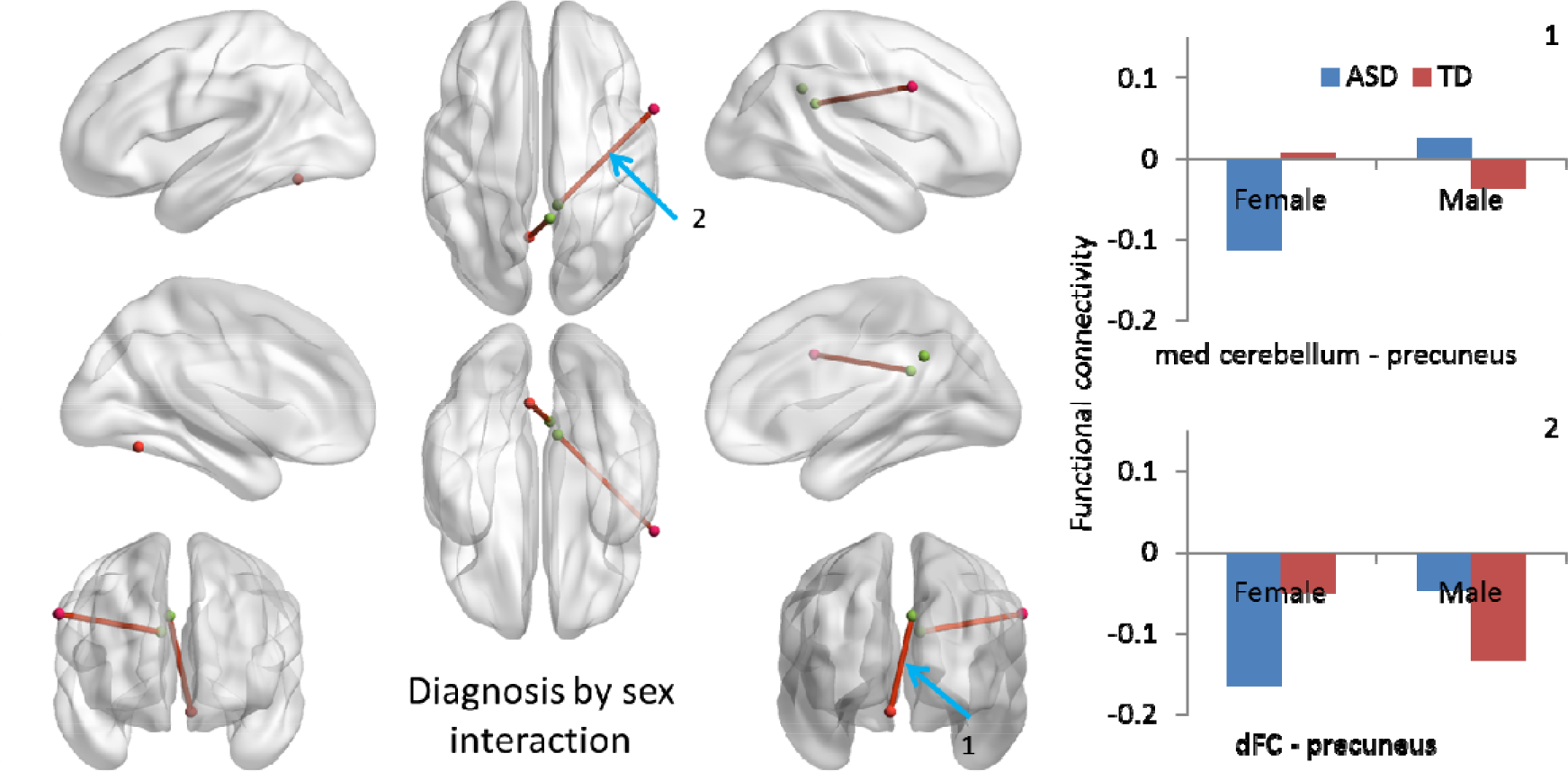
Functional connectivity that showed significant diagnosis by sex interaction at p < 0.0167 after false discovery rate (FDR) correction. Right panels show mean functional connectivity for the four groups of subjects of the corresponding connections.

### 2.3 Functional MRI preprocessing and data analysis

#### 2.3.1 Preprocessing

Functional MRI image preprocessing and analysis were performed using SPM12 (http://www.fil.ion.ucl.ac.uk/spm/) under MATLAB environment (http://www.mathworks.com/). For all the subjects, the first four fMRI images were discarded. The remaining fMRI time-series images were realigned to correct for head motion, and then coregistered to each subject's high resolution anatomical image. The anatomical image of each subject was segmented using the segment routine in SPM. The functional images were then normalized to the standard MNI (Montreal Neurological Institute) space by using the deformation field images obtained from the segmentation step. 24 head motion variables (Friston et al. 1996), five eigenvariates of WM, and five eigenvariates of CSF (Chai et al. 2012) were regressed out for each voxel using a linear regression model. And lastly, the time series from each voxel was band-pass filtered between 0.01 and 0.08 Hz.

Because head motion is a concern in resting-state fMRI (Van Dijk et al. 2012; Power et al. 2012), we performed rigorous steps to estimate and correct for motion in the current analysis. We calculated frame-wise displacement (FD) for translation and rotation separately based on the rigid body head motion estimates that were obtained from the realign procedure. Subjects' data were discarded if the mean FD across any directions exceeded 0.2 mm/deg or maximum FD in any directions exceeded 1.5 mm/deg. We next performed a 2 (diagnosis) by 2 (sex) analysis of variance (ANOVA) on mean FD for each site and for the whole sample. Unfortunately, a significant main effect of diagnosis was observed. Therefore, we included mean FD as covariate in subsequent analysis. A single variable of averaged mean FD in translation and rotation directions was used as a covariate in the group level analysis.

#### 2.3.2 Functional connectivity analysis

We adopted Dosenbach's 160 regions of interest (ROI) to calculate functional connectivity (Dosenbach et al. 2010). Seven ROIs that were located in the cerebellum were discarded, because not all the subjects' fMRI data covered these ROIs. The ROIs were defined as spheres with the center at the provided coordinates (Dosenbach et al. 2010) with a radius of 8 mm. Mean time series was calculated for each ROI in all the subjects. Pearson's correlation coefficients across ROIs were calculated for each subject, resulting in a 153 × 153 correlation matrix for each subject. The correlation coefficients were transformed into Fisher's z scores for further statistical analysis.

Statistical analysis was performed using network-based statistic toolbox (Zalesky et al. 2010). For each connection, a group-level general linear model (GLM) was used to study group differences. The first three columns represented the main effect of diagnosis, the main effect of sex and the diagnosis by sex interaction, respectively. Covariate variables included the effects of age, age2, full scale IQ, mean FD, and five variables representing the site effects. A column of a constant variable was included at the end of the GLM model. There were thus 13 regressors in the GLM model. F statistics was used to examine the effects of diagnosis, sex, and their interaction, separately. False discovery rate (FDR) at 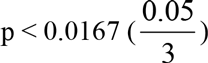 was used to correct for multiple comparisons for all the connections tested. Connectivity results were visualized using BrainMet Viewer (Xia et al. 2013).

For connections that exhibited diagnosis by sex interactions of connectivity, we performed further post-hoc group-wise comparisons by using two sample t-test using the Fisher's z score of connectivity adjusted for age, age2, IQ, head motion and site effects.

## 3 Results

### 3.1 Group effects on IQ and head motion

Even though full scale IQ was matched for each single dataset, when merging the data of all the six datasets, there was a significant effect of diagnosis on full scale IQ after controlling for age, age^2^, and site effects (*F(1,343) = 3.923, p = 0.048*). No significant effect of sex (*F(1, 343) = 0.013, p = 0.908*) and sex by diagnosis interaction (*F(1,343) = 0.921, p = 0.338*) was observed. Therefore, full scale IQ was added in the subsequent analysis as a covariate.

Group level differences were observed in the subject head motion. We observed a significant main effect of diagnosis (*F(1,343) = 12.914, p < 0.001*) and significant effect of sex (*F(1, 343) = 4.324, p = 0.038*) on mean FD after controlling for age, age^2^, and site effects. No significant sex by diagnosis interaction (*F(1,343) = 0.076, p = 0.783*) was found. Therefore, the mean FD was also added as a covariate in subsequent analysis. Mean full scale IQ and FD for each site are listed in Table 1.

**Table 1.** Sample sizes, ages, full scale intelligence quotients (IQ), and mean frame-wise displacement (FD) of included subjects in the current analysis.

|  |  | n |  | Age |  |  |  | IQ |  |  |  | mean FD |  |  |  |
| --- | --- | --- | --- | --- | --- | --- | --- | --- | --- | --- | --- | --- | --- | --- | --- |
|  |  |  |  | ASD |  | TD |  | ASD |  | TD |  | ASD |  | TD |  |
|  |  | ASD | TD | Mean | SD | Mean | SD | Mean | SD | Mean | SD | Mean | SD | Mean | SD |
| PITT | Female | 4 | 2 | 12.6 | 1.0 | 13.6 | 0.5 | 111.5 | 12.3 | 116.5 | 19.1 | 0.075 | 0.018 | 0.075 | 0.022 |
|  | Male | 10 | 10 | 16.0 | 2.0 | 15.3 | 2.7 | 108.4 | 7.0 | 109.8 | 6.3 | 0.077 | 0.029 | 0.078 | 0.021 |
| UM_1 | Female | 7 | 13 | 13.4 | 3.2 | 14.3 | 3.0 | 109.8 | 20.0 | 105.5 | 11.0 | 0.070 | 0.031 | 0.052 | 0.024 |
|  | Male | 20 | 27 | 13.9 | 2.5 | 14.1 | 3.1 | 105.5 | 18.7 | 109.5 | 8.8 | 0.068 | 0.029 | 0.056 | 0.021 |
| YALE | Female | 6 | 7 | 13.9 | 2.6 | 13.9 | 2.6 | 92.0 | 11.4 | 112.3 | 18.3 | 0.057 | 0.014 | 0.057 | 0.021 |
|  | Male | 14 | 17 | 12.7 | 3.3 | 12.6 | 2.9 | 100.9 | 21.7 | 101.4 | 18.4 | 0.066 | 0.024 | 0.060 | 0.031 |
| NYU | Female | 5 | 19 | 13.1 | 2.0 | 11.9 | 2.8 | 101.0 | 23.3 | 113.8 | 15.1 | 0.050 | 0.021 | 0.039 | 0.016 |
|  | Male | 49 | 56 | 11.7 | 3.1 | 12.8 | 3.4 | 108.9 | 16.8 | 112.5 | 13.8 | 0.063 | 0.030 | 0.047 | 0.016 |
| STANFORD | Female | 3 | 4 | 9.4 | 1.3 | 9.0 | 1.0 | 104.3 | 7.0 | 116.0 | 24.7 | 0.055 | 0.022 | 0.055 | 0.005 |
|  | Male | 11 | 9 | 10.3 | 1.6 | 10.4 | 1.9 | 117.8 | 14.5 | 108.6 | 10.3 | 0.080 | 0.020 | 0.071 | 0.037 |
| UCLA_1 | Female | 3 | 4 | 13.6 | 4.2 | 12.8 | 1.1 | 113.7 | 12.6 | 105.5 | 11.1 | 0.082 | 0.054 | 0.045 | 0.025 |
|  | Male | 25 | 22 | 13.7 | 2.6 | 13.6 | 2.1 | 103.8 | 12.3 | 105.0 | 9.4 | 0.060 | 0.022 | 0.051 | 0.022 |
| Total | Female | 28 | 49 | 12.9 | 2.7 | 12.7 | 2.9 | 104.5 | 16.9 | 111.0 | 15.1 | 0.064 | 0.027 | 0.048 | 0.021 |
|  | Male | 129 | 141 | 12.7 | 3.1 | 13.2 | 3.1 | 107.2 | 16.5 | 109.0 | 12.8 | 0.066 | 0.027 | 0.055 | 0.024 |

### 3.2 Resting-state functional connectivity

Mean correlation matrices across 153 ROIs for the four groups are shown in Figure 1. The overall patterns of the matrices were very similar. Because the 153 ROIs were sorted based on their network affiliations defined by Dosenbach et al. (Dosenbach et al. 2010), square like structures along the diagonals are clearly visible.

Functional connectivity between two pairs of ROIs showed significant diagnosis by sex interactions (Figure 2 and Table 2). One was between the precuneus and the medial cerebellum, and the other was between the precuneus and right dorsal frontal cortex. Group mean connectivity strengths of the four groups of subjects for these two connections are shown in the right panels of Figure 2. Post-hoc analysis revealed that for both the connections males with ASD had larger functional connectivity than TD males (connection #1: *t (268) = - 2.1078, SD = 0.1773, p = 0.0360*; connection #2: *t (268) = -2.7972, SD = 0.2030, p = 0.0055*), while females with ASD had smaller functional connectivity than TD females (connection #1: *t(75) = 3.2807, SD = 0.1795, p = 0.0016*; connection #2: *t(75) = 2.6358, SD = 0.2216, p = 0.0102*).

**Table 2.** Resting-state fMRI scanning parameters for the six sites.

|  | Eye status | # of time points | TR (s) | TE (ms) | flip angle (°) | voxel size (mm<sup>3</sup>) |
| --- | --- | --- | --- | --- | --- | --- |
| PITT | Closed | 200 | 1.5 | 20 | 70 | 3.1 x 3.1 x 4 |
| UM_1 | Open | 300 | 2 | 30 | 90 | 3.4 x 3.4 x 3 |
| YALE | Open | 200 | 2 | 25 | 60 | 3.4 x 3.4 x 4 |
| NYU | Open | 180 | 2 | 15 | 90 | 3 x 3 x 4 |
| STANFORD | Closed | 180 | 2 | 30 | 80 | 3.1 x 3.1 x 4.5 |
| UCLA_1 | Open | 120 | 3 | 28 | 90 | 3 x 3 x 4 |

**Table 3.** Connections between regions of interest (ROIs) that had significant effects of diagnosis by sex interaction, and the main effects of diagnosis and sex. Each row represents one connection. X, y, z coordinates were in MNI space (Montreal Neurological Institute). Statistically significant effects were identified by using a threshold of p < 0.0167 after false discovery rate (FDR) correction.

|  | # | ROI 1 |  |  |  | ROI 2 |  |  |  | Effect |
| --- | --- | --- | --- | --- | --- | --- | --- | --- | --- | --- |
|  |  | Label | x | y | z | Label | x | y | z |  |
| Diagnosis by sex interaction | 1 | precuneus | 5 | -50 | 33 | med cerebellum | -6 | -60 | -15 |  |
|  | 2 | dFC | 60 | 8 | 34 | precuneus | 9 | -43 | 25 |  |
| Diagnosis | 1 | post occipital | 13 | -91 | 2 | basal ganglia | 14 | 6 | 7 | T > A |
|  | 2 | precentral gyrus | 58 | -3 | 17 | mid insula | 32 | -12 | 2 | T > A |
|  | 3 | IPS | -32 | -58 | 46 | sup temporal | 42 | -46 | 21 | T > A |
|  | 4 | occipital | 17 | -68 | 20 | temporal | -59 | -47 | 11 | T > A |
|  | 5 | occipital | -29 | -75 | 28 | temporal | -59 | -47 | 11 | T > A |
|  | 6 | mid insula | -36 | -12 | 15 | occipital | 20 | -78 | -2 | T > A |
|  | 7 | precentral gyrus | -54 | -9 | 23 | occipital | 20 | -78 | -2 | T > A |
|  | 8 | frontal | 58 | 11 | 14 | dFC | 60 | 8 | 34 | T > A |
| Sex | 1 | IPS | -36 | -69 | 40 | inf temporal | 52 | -15 | -13 | F > M |
|  | 2 | frontal | 53 | -3 | 32 | occipital | -34 | -60 | -5 | M > F |
|  | 3 | precentral gyrus | 46 | -8 | 24 | occipital | -34 | -60 | -5 | M > F |

T, typically developing
A, Autism spectrum disorder
F, Female
M, Male

Connectivity between eight pairs of ROIs showed significant main effects of diagnosis (Figure 3 and Table 2), with all of them showing smaller connectivity in individuals with ASD than TD individuals. Five connections included one region in the occipital lobe and one other region in the basal ganglia, temporal cortex, middle insula, and precentral gyrus. One connection was between the superior temporal cortex and inferior parietal sulcus. Two other connections were short range connections between the middle insula and precentral gyrus, as well as between the frontal lobe and dorsal frontal cortex. Even though these eight connections did not show a significant diagnosis by sex interaction in connection-wise analysis, the possibility remains that they indeed had small interaction effect but could not survive multiple comparison correction. Therefore, we also checked the diagnosis by sex interaction and the main effect of sex of the eight connections with an uncorrected critical p value of 0.05. No significant diagnosis by sex interaction or main effect of sex was observed even using this liberal threshold of uncorrected p < 0.05.

**Figure 3.**
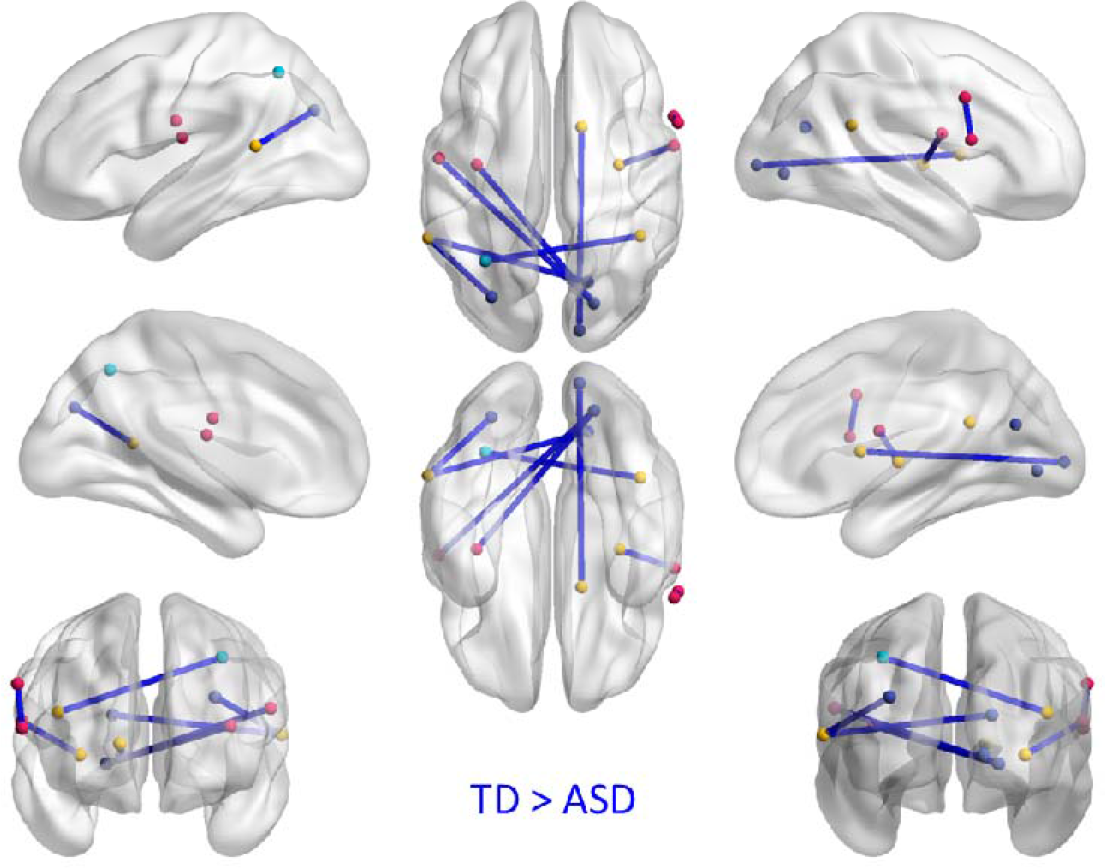
Functional connectivity that showed significant main effect of diagnosis at p < 0.0167 after false discovery rate (FDR) correction. ASD, autism spectrum disorder; TD, typically developed.

Lastly, functional connectivity between three pairs of ROIs showed significant main effects of sex (Figure 4 and Table 2). One connectivity between the left inferior parietal sulcus and right inferior temporal cortex displayed higher connectivity in females than in males. In contrast, the connectivity between the left occipital cortex and right frontal and precentral cortex showed higher connectivity in males than in females.

**Figure 4.**
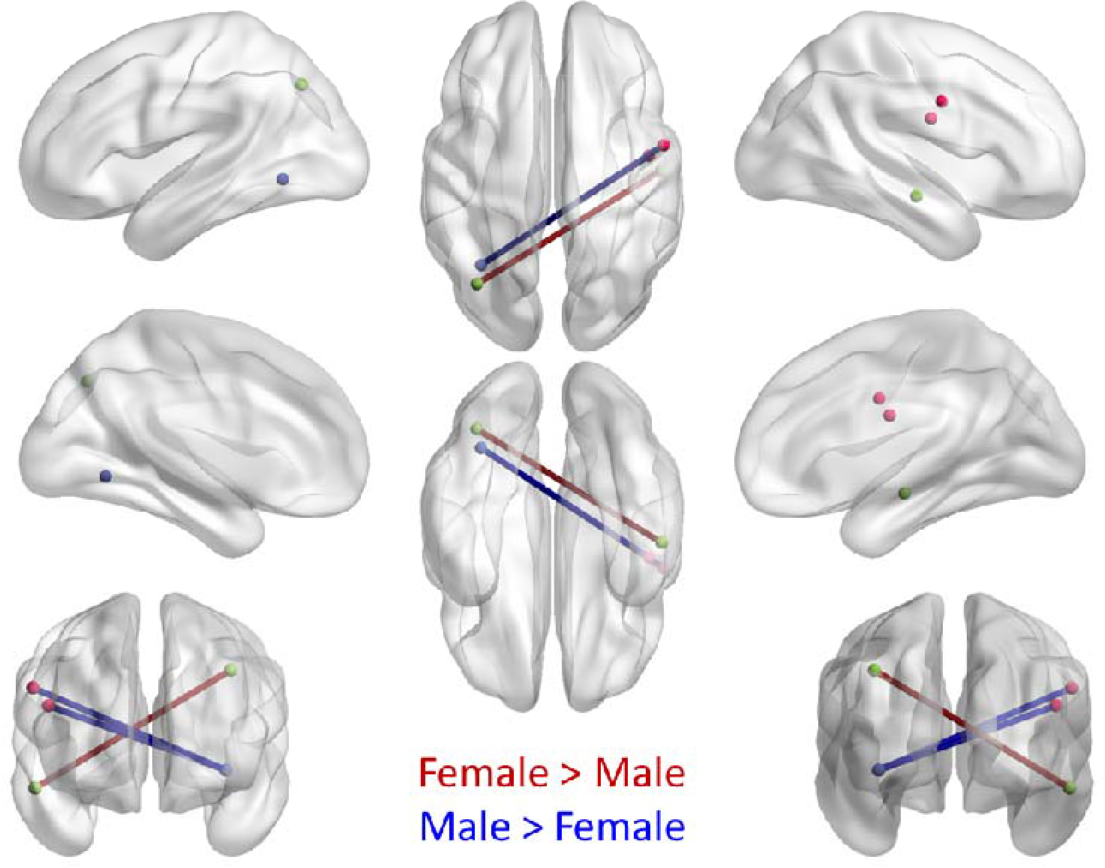
Functional connectivity that showed significant main effect of sex at p < 0.0167 after false discovery rate (FDR) correction.

## 4 Discussion

By aggregating MRI data from a large online data sharing initiative, the current analysis examined sex modulations of resting-state brain connectivity differences in individuals with ASD and TD individuals. Connection-wise analysis identified eight connections that showed only main diagnosis effects, which individuals with ASD had smaller connectivity with TD individuals. More interestingly, we also identified two connections whose connectivity showed a diagnosis by sex interaction. These two connections were between the posterior cingulate cortex and motor region and between the posterior cingulate cortex and cerebellum.

The first observation of the current analysis was that the correlation matrices across ROIs that sampled the whole brain appeared similar among the four groups of subjects. This is in line with the notion that the patterns of resting-state functional connectivity in ASD adults are largely typical (Tyszka et al. 2014). Nevertheless, when performing statistical test on a connection basis, statistically significant differences among groups could still be observed. We observed smaller connectivity in individuals with ASD than TD individuals, mainly between one region from the occipital network and one region from the sensorimotor network or the cinguloopercular network. The smaller connectivity in these connections could be observed in an early study using the same ABIDE dataset, which did not distinguish female and male subjects (Di Martino et al. 2014). However, Di Martino et al. (2014) reported more widely spread smaller and greater connectivity in individuals with ASD. The discrepancies may mainly be due to two factors. First, Di Martino et al. (2014) included more subjects in the analysis, while the current analysis only included datasets with female subjects, and had stringent criteria on IQ and head motion. Secondly, we used a total of 153 ROIs compared with 112 structurally defined ROIs in Di Martino et al. (2014). The total number of multiple comparisons needed to be controlled almost doubled (11,628 vs. 6,216) in the current analysis than in Di Martino et al. (2014). However, the current analysis chose to use functionally defined ROIs (Dosenbach et al. 2010), because the ROIs are more functionally meaningful and have higher spatial resolution. When reducing threshold, similar patterns of greater and smaller connectivity could be observed as Di Martino et al. (2014). Further analysis confirmed that the connectivity showing the main effect of diagnosis did not have diagnosis by sex interaction or main effect of sex, suggesting that these connectivity alterations in ASD are sex-independent. In other words, girls with ASD had similar connectivity reductions in these connections to males with ASD.

More interestingly, we also identified two connections whose connectivity showed a diagnosis by sex interaction. Specifically, TD males showed greater negative connectivity than males with ASD, while females with ASD showed greater negative connectivity than TD females. Such a pattern indicated different alterations in females and males with ASD. Both of the connections involved the precuneus of the default mode network (DMN). The other regions of the connections, which were located in the medial cerebellum and dorsal frontal cortex, may involve in motor functions (Chan et al. 2009). Therefore, the current results may indicate alterations of inhibitory modulation of the DMN on motor functions on motor regions for different sexes. We note that the motor and cerebellar regions are not parts of the typical "task positive network" as defined by Fox et al. (Fox et al. 2005). But these regions still displayed small negative correlations with the precuneus in TD males and females with ASD. The negative correlations between the DMN and task positive networks have been suggested to mediate behavioral variability (Kelly et al. 2008). Behaviorally, females with ASD have been shown to have similarly impaired social cognition than males with ASD, but also relative unimpaired executive function (Lai et al. 2012) and repetitive and restricted behaviors (Supekar and Menon 2015). Therefore, the increased negative correlations between the DMN and motor regions in females with ASD may be a complementary mechanism for them to maintain relatively impaired executive functions. However, we note the difficulties to infer specific functions from resting-state connectivity. More studies using well controlled tasks are needed to study the functions of these connections.

An extreme male brain hypothesis has been proposed to explain sex differences in ASD, stating that the brain of individuals with ASD resembles an extreme form of the male brain (Baron-Cohen et al. 2005; Baron-Cohen 2002). To test this theory, one needs to first observe reliable sex differences in TD individuals, and then check whether individuals with ASD fall toward TD males along the female to male axis. For the two connections that showed diagnosis by sex interactions, TD females had higher functional connectivity than TD males, but males with ASD showed similar pattern as TD females but not TD males. Therefore, the patterns did not agree with the extreme male brain hypothesis. We note that our failing to observe a pattern that favored the extreme male brain theory doesn't preclude that other connectivity had such extreme male brain pattern. Because the current analysis was designed to identify sex-dependent and -independent alterations in ASD, we did not exhaustively test the extreme male brain hypothesis on every connection, which may result in more severe multiple comparison correction problem.

The current study leveraged current large scale data sharing efforts to aggregate a reasonable sample size to study sex differences in ASD. Even though we have carefully controlled for site specific effects, we note the differences in term of methodology used in different sites. Future studies are certainly needed with more methodologically homogeneous sample performed at a single scanner site to replicate the current findings. Another key issue of the current analysis is the unbalanced sample of females and males. The female to male ratio of the ASD subjects in the current analysis is bout 1:4.6, which is comparable to the overall sex ratio of ASD (Baxter et al. 2014; Fombonne 1999). And because the data are already available, there was no reason to remove male subjects to match the sample size of females. Incorporating more subjects is always beneficial for better estimating statistical results. Lastly, we acknowledge that the age range of the current sample is still broad, and covers the transitions to puberty. Our previous analysis on the anatomical data has shown some evidence that the group differences are sensitive to age (Di and Biswal 2015). However, due to the limited number of subjects in the resting-state analysis, it is impossible to examine age effects. Further studies are certainly needed to take into account age effects by using larger sample sizes.

## Acknowledgement

This research was supported by grants from NIH R01AG032088, R01DA038895.

